# Drift-driven microbiome simplification generates reconstructable and ecologically cohesive microbial communities

**DOI:** 10.64898/2026.07.14.738433

**Authors:** Rubén Chaboy-Cansado, Paula Cobeta, Ramón Gallego, Alberto Rastrojo, Daniel Aguirre de Cárcer

## Abstract

Engineering simplified microbial communities that retain function and ecological cohesiveness remains a major challenge because the taxa and interactions required for community establishment are rarely known a priori. Here, we experimentally evaluated drift-driven microbiome simplification as an alternative strategy for generating reduced and reconstructable microbial consortia. Using the tomato rhizosphere as a model system, three source microbial communities were subjected to different dilution bottlenecks and serial propagation. Increasing dilution was the main determinant of community composition and simplification, while repeated passage produced additional reductions in ASV richness, phylogenetic diversity, and evenness. Importantly, diversity loss was not accompanied by a uniform deterioration in bacterial colonization or plant performance, indicating that substantial simplification can occur without a parallel collapse in these system-level properties. Candidate Minimal Microbiome prototypes were selected from endpoint communities, reconstructed as synthetic communities, and their ecological cohesiveness was evaluated through invasion experiments. The reconstructed communities strongly restricted the establishment of the original complex microbial fractions, although invasion success depended markedly on invader identity. The drift-derived communities were at least as resistant to invasion as an independently designed bottom-up synthetic community. Together, these results provide experimental support for drift-driven simplification as a strategy to generate reduced microbial communities whose dominant members can be isolated, reconstructed, and experimentally evaluated. By allowing ecological assembly to generate candidate community configurations before cultivation and reconstruction, this approach provides a complementary route to rational bottom-up design and function-directed top-down microbiome engineering.

## INTRODUCTION

The domestication of microbiomes has emerged as a major goal in modern biology ^1^, with broad applications in agriculture, medicine, and biotechnology ^2^. A central strategy toward this goal is the bottom-up design of synthetic microbial communities by selecting microbial populations expected to collectively perform a desired function. However, this rational engineering approach remains challenging because it requires a detailed understanding of microbial ecology, community assembly, and the functional contributions of individual taxa that is still largely lacking. Recognizing these limitations, Chang et al. (2021) proposed that, rather than attempting to overcome the eco-evolutionary processes governing microbial communities, microbiome engineering could instead harness these processes to generate stable and functional microbial consortia ^1^.

Building on this idea, Talavera-Marcos and Aguirre de Cárcer (2026) recently proposed a microbiome engineering strategy based on the experimental manipulation of ecological drift to isolate and reconstruct cohesive, functionally complete microbial consortia from complex ecosystems ^3^. The approach is founded on the understanding that natural microbiomes are composed of a myriad of local communities associated with different microscale habitat types ^4,5^, each representing a functionally integrated consortium adapted to a particular micro-ecological context. During serial passage under controlled dilution, ecological drift can progressively simplify overall microbiome composition through stochastic extinction and fixation within the metacommunity while preserving microbiome functionality. Rather than recovering intact local communities from the original ecosystem, the process is expected to generate novel but cohesive consortia assembled from populations originating from multiple source communities. The resulting simplified microbiomes are expected to retain the functional potential required to occupy the available niches while containing substantially fewer microbial populations than the original ecosystem, thereby greatly facilitating downstream isolation and microbiome reconstruction. Although various independent studies had provided circumstantial evidence consistent with this concept, Talavera-Marcos and Aguirre de Cárcer (2026) also evaluated drift-driven microbiome simplification using a computational framework reproducing serial dilution–growth dynamics ^3^. Importantly, the simulations were intended to explore the general ecological parameter space governing drift-driven microbiome simplification rather than to reproduce specific microbial ecosystems. Consequently, the results provided general principles for engineering minimal, cohesive and functionally complete microbial consortia while also improving our understanding of analogous processes in heterogeneous natural metacommunities.

The simulations showed that the success of drift-driven microbiome simplification depends on key parameters, particularly the interaction between community size and dilution factor, and the frequency and strength of inter-group biotic interactions. Although the generation of Minimal Microbiomes was predicted to be feasible under a broad range of conditions, the process inevitably involves trade-offs, including the potential loss of functional groups occupying small ecological niches. Nevertheless, the authors also suggested practical strategies to mitigate these limitations, reinforcing the potential of drift-driven simplification as a general approach for engineering low-diversity, functionally cohesive microbial consortia.

However, whether drift-driven simplification can be implemented experimentally in a host-associated microbiome to generate community states that can subsequently be isolated, reconstructed, and retain ecological cohesiveness remains untested. Here, we present an *in vivo* proof of concept of drift-driven microbiome simplification. Using the tomato rhizosphere as a model system, we experimentally manipulated both the source microbial community and the dilution factor applied between serial passages to evaluate how these parameters influence community dynamics throughout the simplification process. We additionally examined whether progressive simplification was associated with changes in bacterial load and plant performance.We further selected candidate Minimal Microbiome prototypes, reconstructed their dominant members as synthetic communities, and used resistance to invasion as an operational test of their ecological cohesiveness and niche occupation. We additionally examined whether progressive simplification was associated with changes in bacterial load and plant performance.

## MATERIALS AND METHODS

### Experimental design

For the initial drift experiment, starting from three initial bacterial communities, tomato plants (*Solanum lycopersicum*) were inoculated with each community and grown for three weeks (Passage 0). After this time, the plants were processed to obtain the three corresponding Rhizosphere Microbial Fractions (hereafter referred to as RMFs). From each RMF obtained at Passage 0, seven dilution levels including the undiluted treatment were prepared to create seven independent experimental dilution trajectories, which marked the beginning of the drift experiment and were used to inoculate the next passage (Passage 1). Each trajectory was established with four biological replicates, each with a technical duplicate. These replicate trajectories were maintained independently throughout the experiment. Thus, for each source community and dilution treatment, four independent experimental trajectories were established. At each passage, each trajectory was represented by two technical replicate plants, of which one was selected for propagation. After three weeks, one technical replicate of each biological replicate within each trajectory was processed to obtain new RMFs, which were then used to inoculate the plants for the subsequent passage. This process was repeated up to six times (Figure 1).

**Figure 1.**
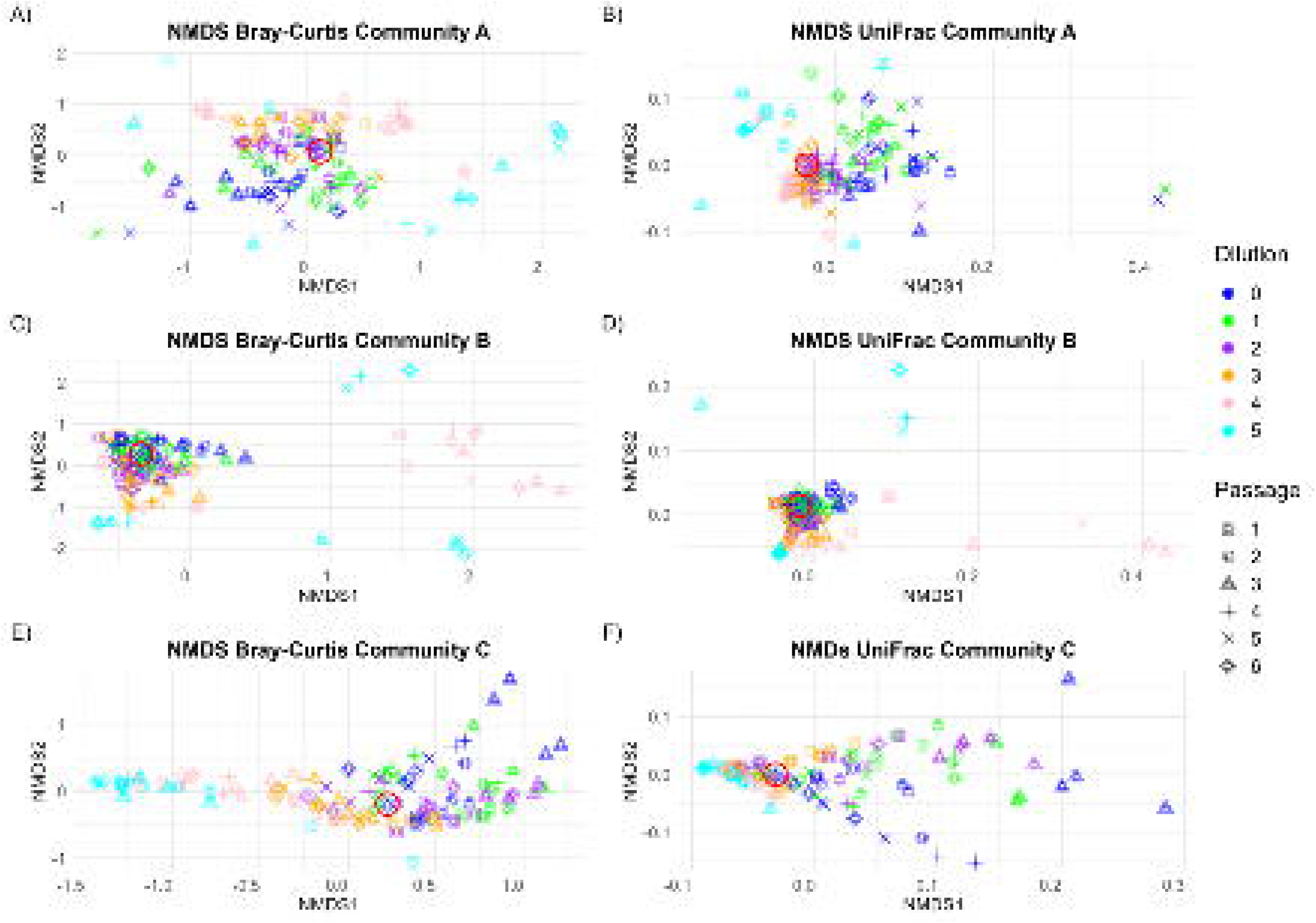
Schematic representation of the procedure used for the production of passage 0 rhizosphere microbial fractions and inoculation of the first two experimental passages. The process was performed using three different soil samples, from which the inocula used to water the plants were prepared.

For the subsequent invasion experiment, we employed four synthetic communities (MMA, MMB, MMC, and PCG) and the three initial microbial fractions (MFA, MFB, and MFC). MMA, MMB, and MMC represent Minimal Microbiome prototypes; one endpoint experimental drift sample for each initial microbial fraction A-C was selected trying to reach an equilibrium between similarity to dilution 0 samples in terms of structure and diversity and bearing a significantly reduced ASV richness. These samples were included in a high-throughput culturomics effort ^6^ that isolated the individual strains responsible for the largest part (97%) of ASVs with relative abundance at least 1% in the samples. For the design of community PCG we instead followed a bottom-up approach and included one representative each for eight of the twelve phylogenetic core groups that were previously detected for the tomato rhizosphere ^7^ and available in our collection. The four synthetic communities were significantly different; MMA was composed of one isolate each affiliated to the genera *Burkholderia*, *Stenotrophomonas*, *Enterobacteriaceae*, and *Herbaspirillum,* two from *Luteibacter*, and three *Pseudomonas*. MMB was composed of one isolate each from *Stenotrophomonas*, *Tatumella*, *Burkholderia*, and *Massilia*. MMC presented one isolate each from *Chitinophaga*, *Herbaspirillum*, *Luteibacter*, *Massilia*, and three *Burkholderia*. PCG was composed of one isolate each from *Pseudarthrobacter*, *Bacillus*, *Variovorax*, *Burkholderia*, *Massilia*, *Shinella*, *Rhizobium*, and *Chryseobacterium*. In this last case, isolates did not all come from the same sample but were retrieved from our general tomato rhizosphere isolates collection. In the invasion experiments, the germinated seed was inoculated with the resident community at the time of planting, then watered with 9 ml of medium bearing the same titer of invading community two weeks later, and the resulting rhizospheres were analyzed two weeks later. With “Control” being a blank inoculation with only watering medium, the first term indicating the resident community and the second term the invading community, the combinations established were: first, resident-only controls (MMA-Control, MMB-Control, MMC-Control and PCG-Control) allowed characterization of rhizosphere communities assembled by each resident synthetic community in the absence of a second inoculum. Second, microbial-fraction treatments (e.g. MMA-MFA, MMA-MFB and MMA-MFC) evaluated the invasion of natural microbial communities introduced into previously assembled synthetic communities. Third, synthetic– synthetic treatments (e.g. MMA-MMB, MMB-MMC) evaluated community cross-invasion. Additional Control-MFA, Control-MFB and Control-MFC treatments were included to characterize rhizosphere communities assembled by each microbial fraction invasion in the absence of a resident synthetic community. After removing poorly performing plants, three to four replicates remained for each combination.

### Preparation of Initial Microbial Fractions and Rhizosphere Microbial Fractions

Microbial Fractions (MFs) had been previously extracted from soils collected in three different locations: the city of Guadalajara (A), the Fuencarral district (B), and the Valdelatas forest (C), the latter two located in Madrid. The first two corresponded to urban recreational orchads, while the third represented a natural forest soil ^8^. For the preparation of the initial Rhizosphere Microbial Fractions (RMFs), 72 tomato plants were used and equally divided into three groups. Each group was inoculated with 100 µL of MFs extracted from one of the three soil types. After three weeks of growth, the 10 best-performing plants per soil were selected and processed, yielding the initial RMFs; from the 20 mL of PBS used to recover the roots in each RMF, up to 15 mL were pooled into sterile 50 mL polypropylene tubes. From each pooled RMF, aliquots were distributed into sterile 1.5 mL tubes. For each RMF, four 1 mL aliquots were used to prepare the inocula for Passage 1, and 750 µL were mixed with 250 µL of sterile 80% glycerol for long-term storage at −80 °C. Serial 1:10 dilutions ranging from 1:1 to 1:10⁶ were prepared from each of the three RMFs obtained at Passage 0, and these were used as inocula for the beginning of the experimental trajectories in Passage 1.

### Plant growth

Previous studies have shown that host genotype can influence rhizosphere microbiome composition ^9,10^. To avoid this potential source of variation, all experiments were performed using seeds from a single genetically homogeneous tomato cultivar, Salinas F1 (CapGen Seeds). Seeds were surface-sterilized and germinated in the dark on water–agar plates ^8^. Germinated seedlings with primary roots approximately 2–3 mm long were transferred individually into sterile 40 mL polypropylene tubes containing 3 g of pre-sterilized vermiculite. Each seedling was immediately inoculated by watering with 15 mL of Gamborg B5 plant culture medium (Duchefa Biochemie) supplemented with the corresponding bacterial inoculum with a titer of 100,000 living celles (MPN). Plants were grown for three weeks in the drift experiments and four in the invasion experiments in sterile greenhouses maintained inside a growth chamber under a 16 h light (24 °C) / 8 h dark (16 °C) photoperiod. Before use, the greenhouses were sterilized by wiping all surfaces with 70% ethanol, sealing ventilation openings with autoclaved cotton, and exposing the interior to ultraviolet light for 5 min.

### Plant Processing

Greenhouses were opened inside a laminar flow hood to retrieve the plant-containing tubes. In the drift experiments, for each plant in passages 1 through 6, the best-performing technical replicate was selected for processing. Selection criteria included the absence of visible fungal growth in the vermiculite and lack of yellowing or dried leaves, or unusually small size.

For all experiments, aerial parts were separated from the roots using sterile forceps. Adhering vermiculite was removed as thoroughly as possible with the same forceps. Shoots were then dried at 65 °C in an oven (JP Selecta, model 200) for 48 hours and weighted using a precision balance. Roots were placed in sterile 40 mL polypropylene tubes containing 2 mL of phosphate-buffered saline (PBS) to extract the rhizospheric microbial fractions (RMFs) and kept cold during the process. Once all roots were transferred to their corresponding tubes, they were vortexed at 1490 rpm for 10 minutes at 4 °C using a Multi Reax vortex mixer (Heidolph).

Following vortexing, 1 mL of each RMF was transferred to a sterile 1.5 mL polypropylene tube. Using an Opentrons OT-2 pipetting robot (Opentrons Labworks Inc.), aliquots were distributed into 0.2 mL polypropylene PCR strip tubes for DNA extraction, storage at −80 °C, microbial titration, and preparation of new inocula. To store the RMFs, 105 µL of each sample was mixed in triplicate with 35 µL of 80% sterile glycerol and frozen at −80 °C. For DNA extraction, 120 µL of each RMF was allocated, and an additional 100 µL from each RMF was used for microbial titration as previously described ^8^.

In the drift experiments, to prepare the inocula for Passage 1, 100 µL from each of the six dilutions of the Passage 0 RMFs, as well as the undiluted RMF, were mixed with 15 mL of sterile irrigation medium. As described previously, each inoculum was used to water four plants, each with one technical replicate. In each of the three greenhouses (one per soil type), two negative control plants were included and irrigated with 15 mL of sterile irrigation medium only. For Passages 2 through 6, RMFs obtained from the previous passage were diluted using the same dilution factor as in the original inoculation. For example, if a plant in Passage 1 had been inoculated with a 1:100 dilution, the RMF recovered from that plant was again diluted 1:100 before being used to inoculate the next plant along with 15 mL of sterile irrigation medium. As in the previous passage, two technical replicate plants per greenhouse were included as negative controls and irrigated only with sterile medium.

### Community profiling

Community DNA was extracted from each RMF using the microvolume alkaline lysis protocol described by Bramucci et al. (2021) ^11^. The V5–V7 region of the bacterial 16S rRNA gene was amplified with primers 799F and 1193R carrying Illumina adapter sequences separated by 0–7 random nucleotides ^11^. PCR amplifications were performed in 25 µL reactions containing 4 µL of template DNA, 0.1 µM of each primer, 0.4 mM dNTPs, and 0.5 U of Q5 High-Fidelity DNA Polymerase (NEB). Cycling conditions consisted of an initial denaturation at 95 °C for 30 s, followed by 25 cycles of 95 °C for 10 s, 55 °C for 30 s, and 72 °C for 30 s, with a final extension at 72 °C for 2 min. Primary PCR products were diluted 1:10 in nuclease-free water before a second 10-cycle PCR using indexed primers carrying Illumina i5/i7 adapter sequences, 10-nt barcodes, and sequencing primer binding sites. Samples within each PCR batch shared one barcode while carrying unique barcodes at the opposite end, allowing unambiguous demultiplexing after sequencing.

Amplicon libraries were examined by agarose gel electrophoresis and pooled at approximately equimolar proportions based on band intensity. Pooled libraries were concentrated using Pellet Paint® NF Co-Precipitant (Merck), size-selected by agarose gel electrophoresis, purified with the Speedtools PCR Clean-Up Kit (Biotools), and sequenced on an Illumina NextSeq platform using a NextSeq1000/2000 P1 flow cell with 2 × 300 bp paired-end reads.

### Sequence processing and analyses

Sequence data were processed with the *DADA2* package in R ^12^ following the standard pipeline for quality filtering, error correction, paired-end merging, chimera removal, taxonomic assignment against the SILVA training database (v123) ^13^, and removal of non-bacterial sequences, including mitochondrial and chloroplast reads. Sequencing depth was standardized by rarefaction using thresholds defined from the natural breaks in sequence-count distributions ^14^ (9,526, 7,895, and 10,961 reads in drift community datasets A, B, and C, respectively, and 63,928 for the invasion community dataset), and samples with fewer reads were discarded. Most samples from the 10⁻⁶ dilution treatment fell below these sequencing-depth thresholds and were therefore excluded; consequently, this dilution treatment was removed from downstream analyses. For phylogeny-based analyses, ASVs were mapped to the Greengenes2 99% representative sequence database and its associated phylogenetic tree ^15^ using BLAST. Faith’s phylogenetic diversity was calculated with the *pd* function from the *picante* package (v1.8.2) ^16^.

## Statistical analyses

For the initial drift experiments, the effects of sampling passage (hereafter, Passage) and dilution level (hereafter, Dilution) on ASV richness (Richness), evenness (Shannon index; Evenness), phylogenetic diversity (Faith’s index; Diversity), bacterial load (Load), and plant dry weight (Weight) were evaluated. For Richness, Evenness, Diversity, and Load, linear mixed models were fitted using the *lmer* function from the *lme4* package (v1.1-37) ^17^, including Passage, Dilution, and their interaction as fixed effects. Trajectory identity was included as a random intercept to account for variation among experimental lines and indirectly among communities. Sum-to-zero contrasts were used for factor coding with the *contr.sum* and *contr.poly* functions in *R*. Load values were log10-transformed prior to analysis to improve normality.

For Weight, mixed models showed singularity due to negligible random-effect variance. Therefore, Weight was analyzed using a linear model with source community identity (Community), Passage, and Dilution as fixed effects. Weight values were converted to milligrams and log10-transformed prior to analysis.

For mixed models, effect significance was assessed using Type III ANOVA with the *anova* function from the *lmerTest* package (v3.1-3) ^18^. For linear models, significance was assessed using Type III ANOVA with the *Anova* function from the *car* package (v3.1-3). Heteroscedasticity in linear models was evaluated using *bptest* from the *lmtest* package (v0.9-40).

To evaluate the effects of Richness and Diversity on Load and Weight, the relationship between both variables was first assessed using Pearson correlation. To obtain the fraction of Diversity independent of Richness, residuals from a linear model of Diversity against Richness were extracted.

The effects of Richness and the Diversity component independent of Richness on Load were analyzed using linear mixed models with additive effects only, including line identity as a random intercept. Load values were log10-transformed prior to analysis. For Weight, singularity again prevented mixed-model fitting, and a linear model including bacterial community, Richness, and independent Diversity as additive fixed effects was fitted instead. Weight values were converted to milligrams and log10-transformed before analysis.

In both cases, effect significance was assessed using Type II ANOVA with the *Anova* function from the *car* package. Heteroscedasticity in linear models was evaluated using *bptest* from the *lmtest* package.

For the ensuing invasion experiments, differences in ASV richness between each treatment and its corresponding control were evaluated using Welch’s two-sample t-tests. P-values were adjusted within each resident community block using the Holm correction for multiple comparisons.

To estimate the establishment of the invasive community, ASVs potentially contributed by each invading community were first identified as those detected in at least one replicate of the corresponding Control_MF treatment. For each resident community, ASVs already detected in the corresponding resident-only controls were removed, yielding a resident-specific set of invasion-marker ASVs for each resident–invader comparison. For each sample, the relative abundance of the invader community was calculated as the summed relative abundance of these candidate ASVs. This value was then multiplied by the total bacterial load of the sample (MPN value) to obtain an estimated absolute invader load. Relative invasion success was calculated for each resident–invader combination as the ratio between the mean estimated invader load in the resident–invader treatment and the mean estimated invader load in the corresponding Control_MF treatment.

### Community composition analyses

Community composition was explored using non-metric multidimensional scaling (NMDS) based on Bray–Curtis and UniFrac distance matrices with the *ordinate* function from the *phyloseq* package (v1.52.0) ^19^. Bray–Curtis dissimilarities were calculated using the *vegdist* function from the *vegan* package (v2.6-10), whereas UniFrac distances were calculated using the *distance* function from *phyloseq*.

To quantify the effects of Passage and Dilution on community composition, partial distance-based redundancy analyses (dbRDA) were performed using Bray–Curtis and UniFrac distance matrices with the *capscale* function from the *vegan* package. In both cases, models were fitted testing the effect of Passage while controlling for Dilution, and vice versa. Effect significance was assessed by permutation tests using the *anova* function, and adjusted R² values were obtained using *RsquareAdj*, both from the *vegan* package.

To evaluate the compositional displacement produced by invasion, for each resident community control, intra-group Bray-Curtis distances were compared to inter-group distances between treatment and control groups using boxplots. Additionally, differences between intra-control and control–treatment distances were assessed using a sample-label permutation test, in which group labels were randomly permuted while keeping the Bray–Curtis distance matrix fixed.

All scripts and 16S rRNA gene dataset are available at https://github.com/microenvgen/Drift and the European Nucleotide Archive (PRJEB114473), respectively. During the preparation of this manuscript, the authors used generative AI tools (ChatGPT, OpenAI) for language editing, translation support, and assistance with code troubleshooting and scripting. All generated suggestions were critically reviewed, validated, and implemented by the authors. The authors take full responsibility for the final content, analyses, and conclusions.

## RESULTS

### Community responses during drift-driven simplification

Across the three experimental communities, rhizosphere microbiomes were dominated by bacterial families commonly associated with this environment, although the dominant families differed among the three inocula. Community A was dominated by Pseudomonadaceae and Xanthomonadaceae, community B by Oxalobacteraceae and Burkholderiaceae, and community C by Burkholderiaceae and Moraxellaceae. In all three cases, marked compositional differences were observed across the initial inoculum dilution treatments. By contrast, community composition remained comparatively stable across serial passages, although some families exhibited gradual changes in relative abundance over time within particular dilution treatments (Figure S1).

To further evaluate differences in community composition, non-metric multidimensional scaling (NMDS) ordinations were performed using Bray–Curtis distances, which are based on community composition, and UniFrac distances, which additionally incorporate the phylogenetic relationships among the ASVs present (Figure 2). Visual inspection of both ordinations suggested that inoculum dilution exerted the strongest effect on community composition, whereas the effect of sampling passage was less evident. To quantify these effects, partial distance-based redundancy analyses (dbRDA) were performed for both distance metrics. The results confirmed the exploratory observations: in all three experimental communities, Dilution consistently explained a larger proportion of the variation in community composition than Passage, although Passage also accounted for a small but significant fraction of the variation for both Bray–Curtis and UniFrac distances (Table 1).

**Figure 2.**
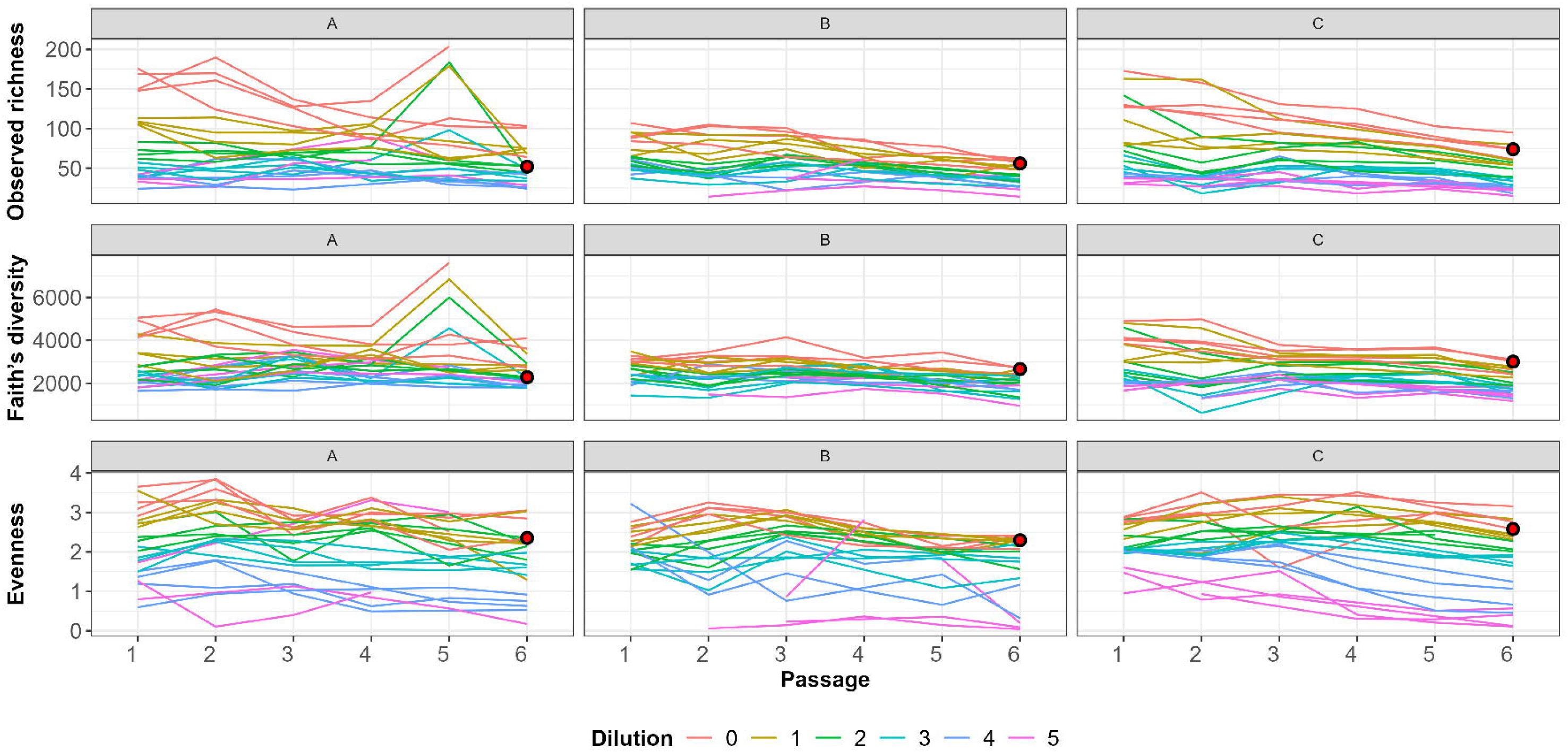
Non-metric multidimensional scaling (NMDS) ordinations based on Bray–Curtis (left) and UniFrac (right) distances of the microbial communities from samples inoculated with communities A–C (top to bottom). Sampling passage is represented by point shape and dilution treatment by color. The endpoint sample selected from each source community as a candidate Minimal Microbiome prototype is highlighted with a red circle.

**Table 1.** Results of the partial distance-based redundancy analyses (dbRDA) based on Bray– Curtis and UniFrac dissimilarities. The percentages represent the proportion of variance explained, expressed as adjusted R². The effect of each variable was estimated while controlling for the other variable, and statistical significance was assessed using 999 permutations.

| Inoculated community | Variable | Explained variance (Bray–Curtis dissimilarities) | Explained variance (UniFrac dissimilarities) | P value |
| --- | --- | --- | --- | --- |
| A | Dilution | 9.81 % | 22.33 % | p < 0.001 |
|  | Passage | 3.36 % | 1.86 % | p < 0.001 |
| B | Dilution | 18.13 % | 15.17 % | p < 0.001 |
|  | Passage | 1.25 % | 0.38 % | p < 0.001 |
| C | Dilution | 30.10 % | 30.83 % | p < 0.001 |
|  | Passage | 9.27 % | 5.03 % | p < 0.001 |

ASV richness decreased significantly with both sampling passage and increasing dilution (Figure 3, top). However, the significant Passage × Dilution interaction indicated that the decline in richness across successive passages was progressively less pronounced as dilution increased (Passage: F₁,₂₉₁.₅₈ = 92.30, P < 0.001; Dilution: F₁,₇₂.₃₉ = 149.17, P < 0.001; Passage × Dilution: F₁,₂₉₅.₅₄ = 56.24, P < 0.001).

**Figure 3.**
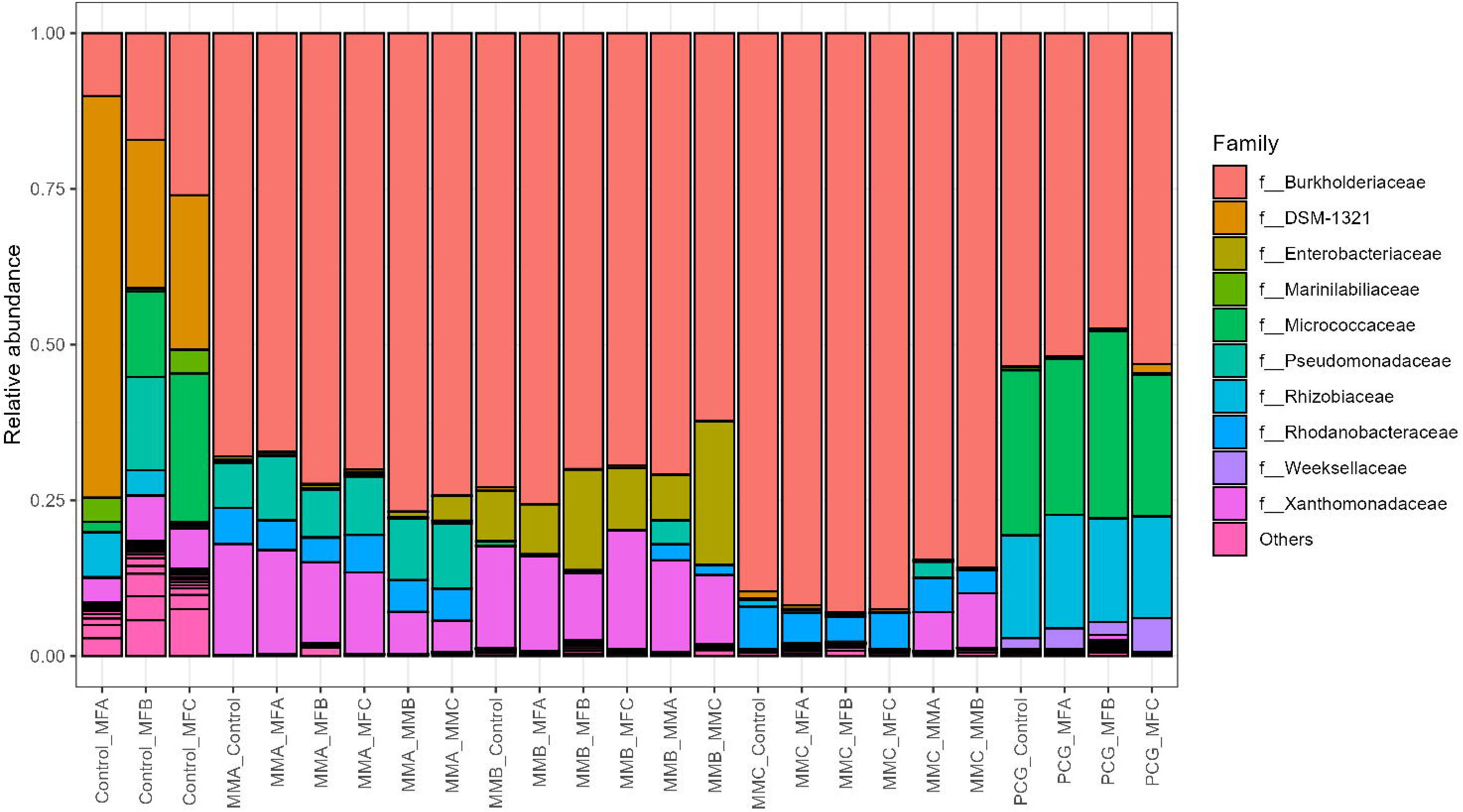
Distribution of Richness, Diversity, and Evenness (top to bottom) as a function of dilution and sampling passage. Each column corresponds to one inoculated community (A–C, left to right). Dilution treatments are represented by different colors, while individual experimental trajectories are shown as separate lines. The endpoint sample selected from each source community as a candidate Minimal Microbiome prototype is highlighted with a red circle.

Similarly, phylogenetic diversity (Faith’s index) decreased significantly with both sampling passage and increasing dilution (Figure 3, center). However, a significant Passage × Dilution interaction indicated that the decline in phylogenetic diversity across successive passages became progressively less pronounced at higher dilution levels (Passage: F₁,₂₉₂.₆₅ = 22.90, *P* < 0.001; Dilution: F₁,₇₃.₀₈ = 85.69, *P* < 0.001; Passage × Dilution: F₁,₂₉₇.₀₉ = 12.18, *P* < 0.001).

Community evenness (Shannon index) decreased significantly with both sampling passage and increasing dilution (Figure 3, bottom). A significant Passage × Dilution interaction further indicated that the decline in evenness across successive passages became more pronounced at higher dilution levels (Passage: F₁,₂₈₅.₉₁ = 49.21, *P* < 0.001; Dilution: F₁,₆₅.₃₅ = 262.55, *P* < 0.001; Passage × Dilution: F₁,₂₉₁.₇₁ = 4.08, *P* < 0.05).

### Experimental effects on bacterial load and plant weight

Bacterial load showed no significant effect of sampling passage, but a marginal tendency to decrease with increasing dilution (Figure S2). However, a significant Passage × Dilution interaction indicated that the negative effect of dilution on bacterial load became progressively weaker across successive passages (Passage: *P* = 0.18; Dilution: F₁,₆₆.₉₃ = 3.18, *P* = 0.07; Passage × Dilution: F₁,₂₂₇.₀₀ = 4.70, *P* < 0.05).

Plant dry weight decreased significantly with sampling passage but increased with increasing dilution (Figure S3) (Passage: F₁,₃₄₉ = 10.78, *P* < 0.05; Dilution: F₁,₃₄₉ = 26.82, *P* < 0.001).

Because ASV richness and phylogenetic diversity were highly correlated (Pearson’s *r* = 0.90, *P* < 0.001), the effects of Richness and Diversity on bacterial load and plant dry weight were analyzed using both Richness and the fraction of Diversity independent of Richness. Both Richness and the Diversity component independent of Richness were negatively associated with bacterial load (Richness: χ² = 7.55, *P* < 0.05; independent Diversity: χ² = 4.40, *P* < 0.05). In contrast, plant dry weight was negatively associated with Richness but showed no significant relationship with the Diversity component independent of Richness (Richness: F₁,₃₅₈ = 19.0, *P* < 0.001; independent Diversity: *P* = 0.80).

### Resistance to invasion by reconstructed Minimal Microbiomes

Based on the combined consideration of reduced ASV richness and retention of some compositional resemblance to the corresponding source community, one endpoint community from each source was selected as a candidate Minimal Microbiome prototype (Figures 2–3). Its dominant bacterial populations were isolated by high-throughput culturomics and used to reconstruct three synthetic communities (MMA, MMB, and MMC). As a reference, a fourth synthetic community (PCG) was independently assembled using a bottom-up strategy based on previously described phylogenetic core groups of the tomato rhizosphere. To evaluate the ecological cohesiveness of these reconstructed communities, they were subjected to microbial invasion experiments. Under this framework, resistance to invasion was used as an operational indicator of the extent to which the resident communities occupied the ecological opportunities available in the rhizosphere under the tested conditions.

Initial exploration of the invasion dynamics (Figures 4-5) suggested that the resident communities displayed substantial resistance to invasion. To quantify this effect, we compared the within-treatment dissimilarity of each resident-only control with the dissimilarity between the corresponding control and its invasion treatments (Figure 6). Significant changes were detected only in a subset of invasions involving other Minimal Microbiomes, whereas none of the invasions performed with the complex initial microbial fractions produced significant shifts in community composition.

**Figure 4.**
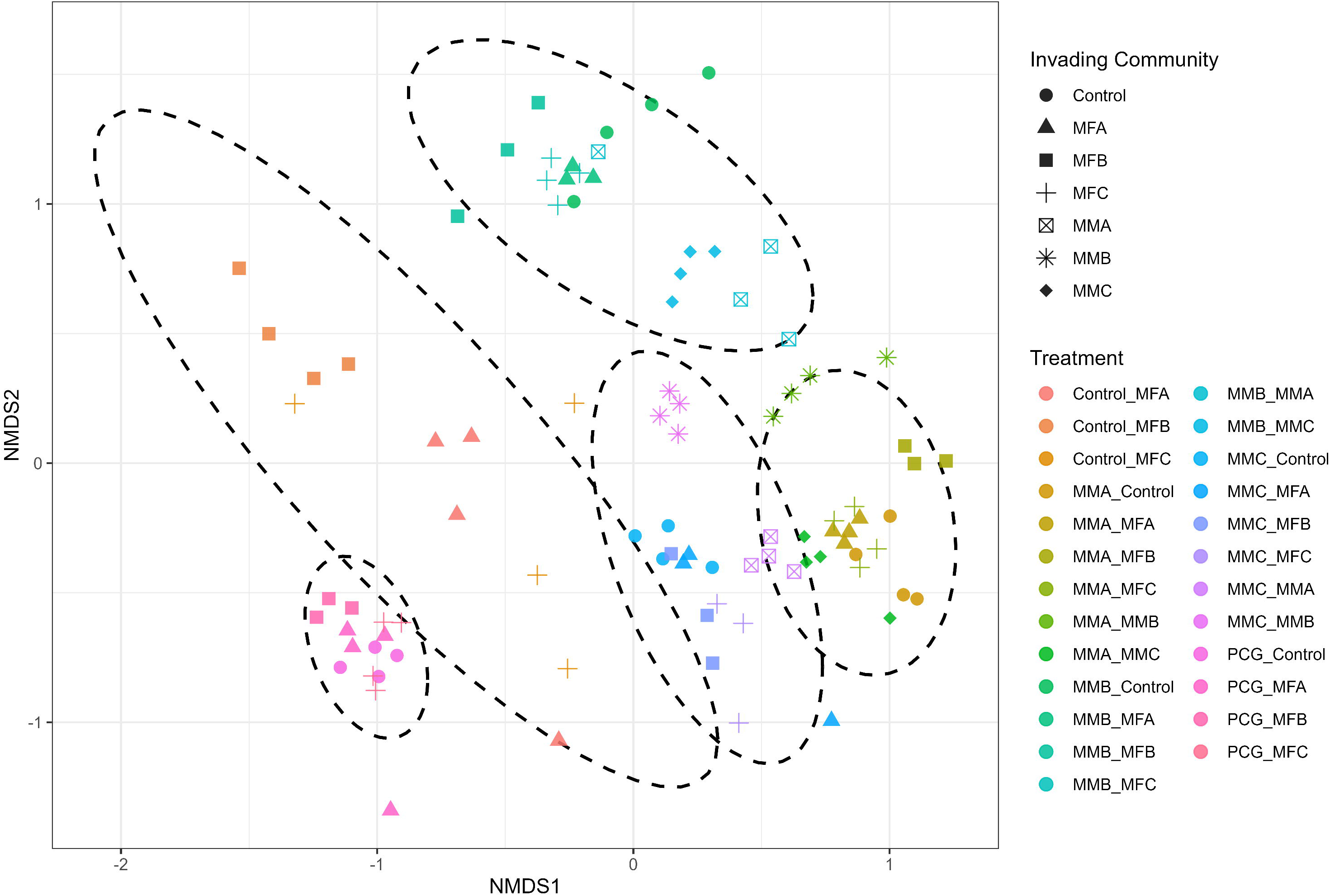
Relative abundance of the ten most abundant bacterial families across the dataset in the different invasion treatments. Each bar represents the mean relative abundance of each family across biological replicates for the corresponding treatment. Treatment names indicate the resident community as the first term and the invading community as the second term. “Control” denotes a blank inoculation containing only the watering medium.

**Figure 5.**
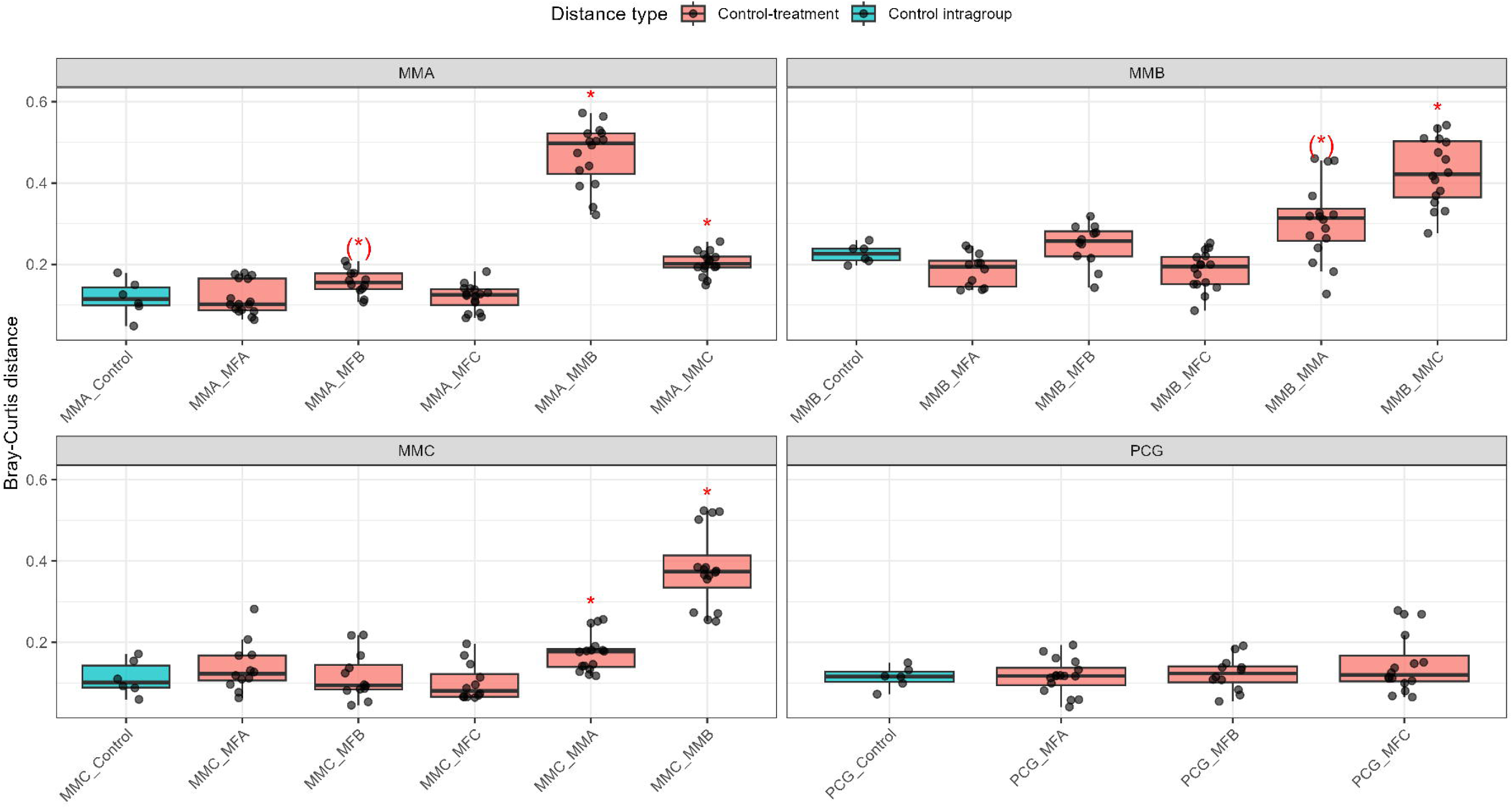
Non-metric multidimensional scaling (NMDS) ordination based on Bray–Curtis dissimilarities among rhizosphere communities from the invasion assays. Each point represents an individual sample. Colors identify the different treatments, whereas point shapes indicate the invading community. Treatment names indicate the resident community as the first term and the invading community as the second term, with “Control” denoting a blank inoculation containing only the watering medium. Ellipses enclose samples sharing the same resident community, irrespective of the invading community.

**Figure 6.**
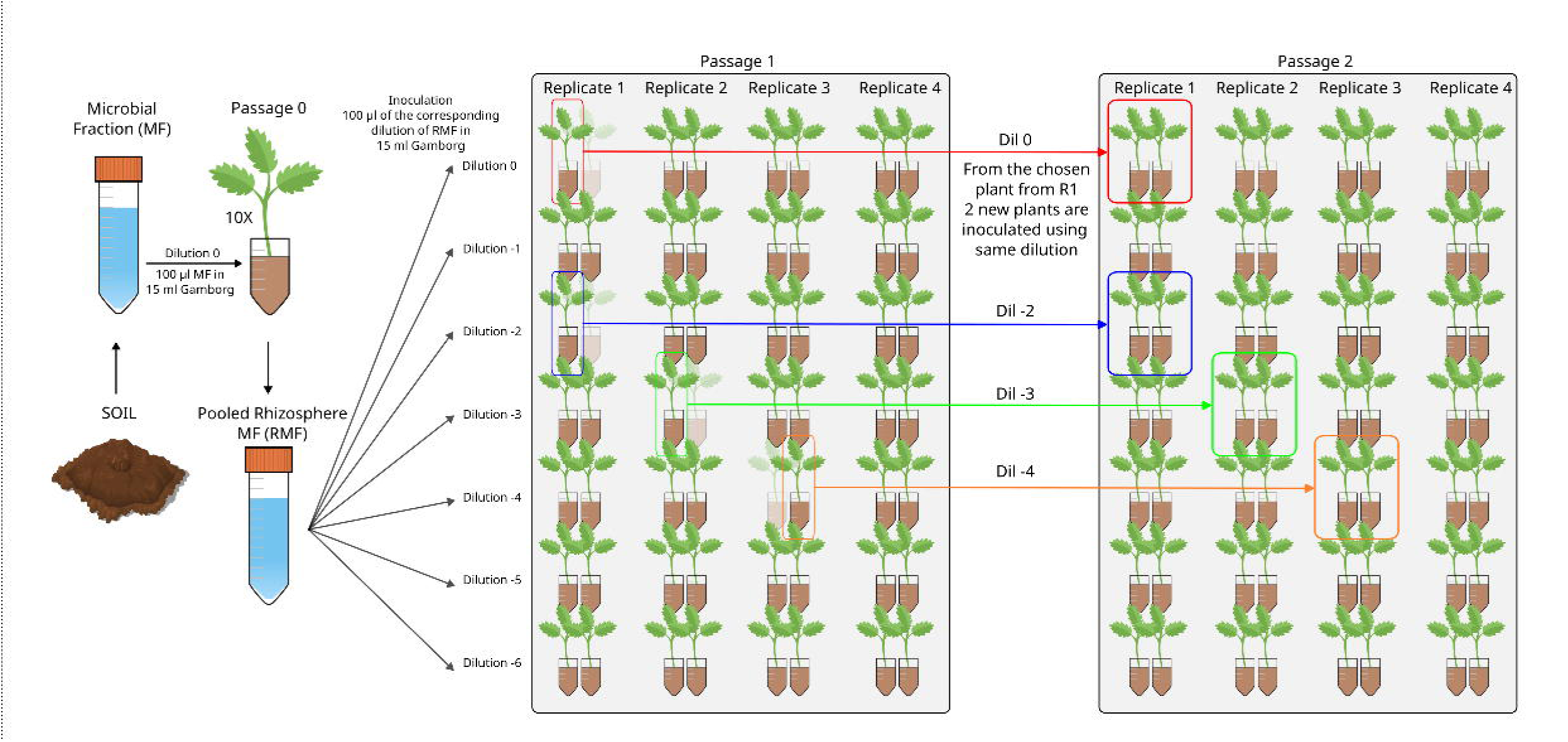
Bray–Curtis distances comparing resident-only controls with their corresponding invasion treatments. For each resident synthetic community, the first boxplot represents the distribution of Bray–Curtis distances among biological replicates of the resident-only control (Control intragroup), whereas the remaining boxplots show the Bray–Curtis distances between control replicates and the corresponding invasion treatment (Control–treatment). Red asterisks indicate significant increases in Bray–Curtis distance relative to the corresponding resident-only control based on permutation tests (999 permutations; unadjusted *P* values): (*) *P* ≤ 0.10, * *P* ≤ 0.05.

It is important to note that the Control–MF treatments, in which the microbial fractions were introduced only two weeks after planting, displayed substantially lower bacterial loads than the remaining treatments, which had a resident community in the planting substrate from the time of planting. Mean bacterial loads were 4.42 × 10⁶ (± 2.16 × 10⁷) MPN units in the Control–MF treatments and 4.82 × 10⁷ (± 1.85 × 10⁸) units in the remaining treatments. This difference was consistent with the shorter colonization period available following late inoculation. Importantly, negative control plants that received no microbial inoculum remained below the detection limit, indicating negligible background colonization. Therefore, the invasion results should be interpreted considering that the invading microbial fractions had less time to establish and proliferate, and this asymmetry is inherent to the experimental design required to assess invasion resistance.

Taking this consideration into account, we next focused on the ability of the invading communities to establish within resident microbiomes. We first examined differences in the detection of bacterial families and ASVs between each invasion treatment and its corresponding resident-only control. We then focused on invasion-marker ASVs, defined for each resident– invader comparison as ASVs detected in at least one biological replicate of the corresponding Control–MF treatment but absent from all biological replicates of the corresponding MM– Control. For each sample, the relative abundances of these invasion-marker ASVs were summed and multiplied by the total bacterial load of that sample to estimate the bacterial load attributable to the invading community, thereby accounting for differences in total bacterial load among treatments. Relative invasion success was then expressed as a percentage of the corresponding Control–MF treatment. Thus, a value of 100% indicates that the invading community reached the same estimated bacterial load as when inoculated in the absence of a resident community, values below 100% indicate reduced establishment in the presence of the resident community, and values above 100% indicate greater establishment than in the corresponding Control–MF treatment.

With respect to total ASV richness, significant increases were observed primarily following MFB invasion of PCG and MMA, with only one additional significant increase detected in another treatment (Table 2). Because the subsequent analysis aimed to quantify establishment of the complex microbial fractions, relative invasion success was calculated specifically for MFA–MFC invasions. Consistent with this pattern, MFB displayed substantially higher relative invasion success than either MFA or MFC across all resident communities (10.9–37.6% versus 0.6–9.3%; Table 3). Overall, the synthetic communities exhibited strong resistance to invasion, with MMA– MFB representing the only treatment in which the invading community achieved a substantial level of establishment (37.6%).

**Table 2.** ASV and family richness by treatment. Values indicate mean ± SD. ASV and family richness were calculated at three thresholds (0.1%, 0.01%, and 0%). Asterisks indicate significant differences in ASV richness between treatment and relevant MM_Control.

| Treatment | ASVs 0.1% | Families 0.1% | ASVs 0.01% | Families 0.01% | ASVs 0% | Families 0% |
| --- | --- | --- | --- | --- | --- | --- |
| Control MFA | 28.0 ± 8.9 | 14.8 ± 4.3 | 60.2 ± 11.7 | 21.5 ± 2.6 | 71.5 ± 10.9 | 22.8 ± 2.6 |
| Control_MFB | 47.5 ± 10.9 | 20.0 ± 0.8 | 108.0 ± 29.6 | 26.8 ± 3.1 | 141.5 ± 48.9 | 30.0 ± 6.2 |
| Control_MFC | 37.0 ± 10.7 | 16.2 ± 2.6 | 87.5 ± 23.6 | 24.5 ± 3.0 | 110.5 ± 32.5 | 25.8 ± 2.9 |
| MMA Control | 18.2 ± 2.6 | 7.0 ± 1.2 | 28.8 ± 2.8 | 10.2 ± 1.3 | 34.2 ± 5.0 | 11.2 ± 2.2 |
| MMA MFA | 16.2 ± 1.9 | 7.2 ± 1.0 | 29.5 ± 5.1 | 12.8 ± 3.2 | 46.8 ± 5.9 | 16.5 ± 3.7 |
| MMA MFB | 21.7 ± 3.2 | 10.3 ± 0.6 | 61.0 ± 3.5* | 19.3 ± 0.6 | 90.7 ± 2.5* | 22.7 ± 2.5 |
| MMA MFC | 16.8 ± 4.0 | 8.5 ± 0.6 | 36.0 ± 6.2 | 13.5 ± 2.9 | 52.8 ± 7.0* | 18.5 ± 5.1 |
| MMA MMB | 20.2 ± 2.1 | 6.2 ± 1.0 | 35.0 ± 5.8 | 10.0 ± 2.6 | 44.2 ± 8.5 | 13.0 ± 2.2 |
| MMA MMC | 19.2 ± 4.3 | 7.2 ± 1.5 | 31.8 ± 2.1 | 11.0 ± 0.0 | 44.5 ± 8.5 | 14.5 ± 2.6 |
| MMB Control | 21.5 ± 3.4 | 6.5 ± 1.3 | 36.8 ± 8.3 | 12.2 ± 3.9 | 51.5 ± 16.1 | 15.2 ± 6.1 |
| MMB MFA | 19.3 ± 4.5 | 7.0 ± 2.6 | 39.0 ± 6.1 | 11.3 ± 2.5 | 58.0 ± 14.5 | 16.0 ± 3.0 |
| MMB_MFB | 23.7 ± 2.1 | 10.7 ± 2.3 | 63.0 ± 16.5 | 19.0 ± 1.0 | 104.7 ± 27.6 | 25.3 ± 4.5 |
| MMB MFC | 17.5 ± 6.6 | 7.8 ± 2.2 | 36.2 ± 11.2 | 11.8 ± 2.1 | 56.5 ± 17.3 | 16.5 ± 3.4 |
| MMB MMA | 28.2 ± 4.5 | 7.8 ± 0.5 | 46.2 ± 6.1 | 11.8 ± 1.5 | 65.2 ± 9.8 | 15.8 ± 1.7 |
| MMB MMC | 28.0 ± 3.2 | 8.5 ± 0.6 | 49.2 ± 7.8 | 13.8 ± 1.0 | 69.0 ± 7.2 | 17.0 ± 3.4 |
| MMC Control | 21.2 ± 3.3 | 7.2 ± 1.7 | 37.0 ± 7.0 | 12.5 ± 2.6 | 56.8 ± 9.2 | 18.2 ± 3.3 |
| MMC MFA | 21.3 ± 1.5 | 7.3 ± 2.1 | 35.0 ± 5.6 | 11.7 ± 2.5 | 42.7 ± 11.7 | 13.7 ± 3.2 |
| MMC MFB | 21.0 ± 4.6 | 9.0 ± 2.0 | 49.3 ± 7.5 | 16.3 ± 4.0 | 85.3 ± 10.4 | 22.3 ± 6.8 |
| MMC MFC | 16.0 ± 1.7 | 5.3 ± 0.6 | 29.0 ± 2.6 | 10.3 ± 0.6 | 43.7 ± 7.5 | 15.3 ± 2.1 |
| MMC MMA | 23.8 ± 1.9 | 8.2 ± 1.5 | 37.8 ± 3.9 | 12.0 ± 2.2 | 50.8 ± 3.8 | 15.8 ± 2.2 |
| MMC MMB | 24.8 ± 3.0 | 7.8 ± 1.5 | 43.5 ± 5.2 | 11.2 ± 1.9 | 61.5 ± 7.5 | 14.8 ± 3.1 |
| PCG Control | 18.0 ± 1.4 | 8.8 ± 1.3 | 39.2 ± 9.3 | 15.0 ± 4.2 | 47.0 ± 9.1 | 17.0 ± 4.2 |
| PCG MFA | 16.2 ± 3.4 | 8.8 ± 1.0 | 40.0 ± 5.6 | 17.2 ± 3.3 | 64.5 ± 15.3 | 22.5 ± 1.3 |
| PCG MFB | 23.7 ± 1.5* | 13.7 ± 0.6 | 55.0 ± 1.7 | 21.7 ± 2.5 | 88.7 ± 5.0* | 24.0 ± 3.5 |
| PCG MFC | 15.0 ± 4.2 | 7.8 ± 2.2 | 39.0 ± 5.4 | 14.2 ± 1.3 | 51.2 ± 3.3 | 17.0 ± 1.4 |

**Table 3.** Relative invasion success of the initial microbial fractions (MFA–MFC) into each resident synthetic community.

| **Treatment** | **Relative invasion success** |
| --- | --- |
| MMA_MFA | 3.08 % |
| MMB_MFA | 1.40 % |
| MMC_MFA | 1.27 % |
| PCG_MFA | 2.38 % |
| MMA_MFB | 37.60 % |
| MMB_MFB | 10.87 % |
| MMC_MFB | 11.30 % |
| PCG_MFB | 19.27 % |
| MMA_MFC | 9.27 % |
| MMB_MFC | 0.77 % |
| MMC_MFC | 0.56 % |
| PCG_MFC | 3.04 % |

## DISCUSSION

In this study, we experimentally explored whether drift-driven community simplification could be used to generate reduced rhizosphere microbiomes that retained the capacity to assemble into ecologically cohesive communities. To this end, three initial tomato rhizosphere microbial fractions were subjected to different dilution bottlenecks and serial propagation, generating progressively simplified communities from which candidate Minimal Microbiome prototypes were selected. The dominant bacterial populations of these prototypes were subsequently recovered through an independent in-house culturomics effort and used here to reconstruct three synthetic communities, whose ecological cohesiveness was evaluated through invasion experiments and compared with a reference synthetic community designed independently through a bottom-up approach. Because reconstruction was based on the dominant populations recovered through cultivation, the synthetic communities should be regarded as culturable approximations of the selected endpoint states rather than exact replicas of their full original composition. Inoculum dilution was the main determinant of both community composition and the extent of microbiome simplification, whereas serial passage produced comparatively smaller compositional changes. Nevertheless, both increasing dilution and successive passages progressively reduced ASV richness, phylogenetic diversity and community evenness, consistent with the cumulative loss of microbial populations expected under repeated population bottlenecks. From the resulting communities, candidate Minimal Microbiome prototypes were qualitatively selected by seeking a practical balance between sufficiently low richness to make isolation and reconstruction feasible and the retention of some compositional resemblance to their respective source communities. Rather than designing these communities a priori, drift-driven simplification therefore allowed the ecological assembly process itself to generate simplified candidate configurations, which were subsequently contrasted with PCG, an independently designed bottom-up community based on phylogenetic core groups of the tomato rhizosphere. Importantly, drift-driven diversity loss was not accompanied by a uniform deterioration in either bacterial colonization or plant growth. Bacterial load did not decline significantly across passages, although it showed a marginal decrease with increasing dilution, while plant dry weight decreased across passages but increased with dilution. Moreover, both richness and phylogenetic diversity independent of richness were negatively associated with bacterial load, and richness was negatively associated with plant dry weight. Together, these results do not support a simple positive relationship between microbial diversity and community performance in this system. Rather, the consequences of simplification depended on how it was generated, with dilution and repeated passage producing partly contrasting effects. These observations suggest that substantial diversity loss can occur without necessarily causing a parallel collapse in rhizosphere colonization or plant performance, although they do not constitute a direct functional validation of the subsequently reconstructed Minimal Microbiomes.

The ability of the reconstructed communities to resist invasion was used as an operational measure of ecological cohesiveness and niche occupation. Established microbial communities can restrict subsequent colonists through resource depletion and niche pre-emption, occupation of physical space, environmental modification, antagonistic interactions, or host-mediated feedbacks. Thus, although invasion resistance cannot demonstrate that all functions of the original microbiome have been retained, strong restriction of incoming populations indicates that the resident consortium collectively exploits a substantial fraction of the ecological opportunities available under the tested conditions.

Overall, the reconstructed communities strongly restricted the establishment of the complex microbial fractions, although invasion success was clearly dependent on invader identity. Interestingly, the few significant global compositional shifts were observed in cross-invasions between reconstructed Minimal Microbiomes rather than following invasion by the more complex microbial fractions, further indicating that invasion outcomes depended on community identity rather than simply on invader complexity. MFB consistently established more successfully than MFA or MFC across all resident communities, reaching 10.9–37.6% of its establishment in the corresponding resident-free control, compared with only 0.6–9.3% for MFA and MFC. This pattern indicates that invasion resistance was not an absolute property of the resident community but depended strongly on the identity of the invading community. Notably, MFB originated from a soil with a long history of tomato cultivation, whereas MFA originated from a mixed vegetable garden and MFC from a pine forest. Previous community-coalescence experiments have shown that historical exposure to similar plant hosts can generate specialized microbial communities with a strong competitive advantage during community mixing ^20^, making the consistently greater establishment of MFB in the tomato rhizosphere biologically plausible. Nevertheless, the present experiment cannot determine whether its greater invasiveness resulted from previous adaptation to the tomato rhizosphere, intrinsic colonization traits of its constituent populations, antagonistic interactions with resident taxa, or access to resources insufficiently exploited by the resident communities. The particularly high establishment observed for MMA–MFB (37.6%) further suggests a resident–invader-specific opportunity for establishment; however, because MFB was the most successful invader across all four resident communities, this result cannot be attributed solely to incomplete niche occupation by MMA.

The drift-derived Minimal Microbiomes were at least as resistant to invasion as PCG, which was independently constructed through an ecologically informed bottom-up approach. This indicates that both strategies can generate cohesive rhizosphere consortia. The main advantage of drift-driven simplification may therefore lie not in necessarily producing more resistant communities, but in generating multiple simplified, self-assembled candidate configurations without requiring prior identification of the taxa or interactions needed for community establishment. Moreover, because simplification occurs before isolation, it significantly facilitates the recovery and subsequent reconstruction of the community members.

Our approach can be placed within the broader framework of top-down microbiome engineering, here understood as strategies that start from a complex community or metacommunity and modify its composition or function through environmental selection, serial propagation, community-level selection, bottlenecks, or other ecological perturbations, rather than initially designing the community through the rational combination of isolated strains. This general framework has been extensively discussed in conceptual and review studies covering approaches such as enrichment, artificial community selection, directed evolution, and microbiome breeding ^21–25^. The potential and limitations of these approaches have also been explored through theoretical and simulation-based studies ^1,26,27^. Importantly, top-down strategies have also been implemented experimentally, including selection of whole microbial ecosystems for emergent functions, host-mediated selection based on plant phenotypes ^28,29^, and the simplification of enriched functional communities through dilution-to-extinction or related reductive approaches ^30,31^. Together, these studies illustrate that top-down microbiome engineering encompasses a diverse set of strategies in which community-level variation is generated and subsequently filtered or selected, either toward a predefined function or toward reduced community configurations with desirable properties.

This study provides experimental support for the use of drift-driven simplification as a strategy to generate reduced and reconstructable microbiomes, extending previous conceptual and simulation-based work by implementing the complete workflow in a host-associated microbial system. Consistent with the previous simulations, dilution intensity strongly determined the extent of simplification, and repeated propagation produced additional diversity loss. The present study further extends those predictions by showing that selected endpoint states can be recovered through cultivation, reconstructed, and experimentally tested as synthetic communities. Together, our results support the ecological cohesiveness of the reconstructed communities under the tested conditions, while functional properties, temporal stability, resilience to perturbation and effects on the host remain to be independently evaluated.

The experimental design also defines the scope of this conclusion. Resident communities were introduced at planting, whereas invading communities were introduced two weeks later; therefore, the assay evaluates invasion into an older, already established community rather than competition between communities of equivalent age and biomass. The observed resistance may consequently reflect not only niche occupation, but also priority effects arising from previous resource use, spatial occupation, environmental modification, antagonistic interactions or host-mediated feedbacks. Relative invasion success was normalized against the corresponding late-inoculation Control–MF, thereby benchmarking each invader against its establishment under the same late-inoculation schedule in the absence of a resident community. Nevertheless, the assay specifically measures the capacity of a late-arriving community to establish within a pre-assembled resident microbiome. This priority-mediated resistance is itself relevant for preventive microbiome applications, in which successful establishment before exposure to incoming microorganisms may constitute a desirable property.

Overall, our results show that experimental manipulation of drift can generate simplified microbiomes whose dominant members can be isolated and reconstructed into ecologically cohesive consortia, which can subsequently be screened for specific functional properties when required. This approach provides a complementary route to rational bottom-up design and function-directed top-down selection, while successful invaders may additionally identify opportunities for subsequent community refinement.

## Notes

https://github.com/microenvgen/Drift

### Competing Interest Statement

The authors have declared no competing interest.

