## Supplementary for "Drift-driven microbiome simplification generates reconstructable and ecologically cohesive microbial communities"

^1^Microbial and Environmental Genomics Group, Departamento de Biología, Universidad Autónoma de Madrid, 28049, Spain.

^#^Equal contribution.

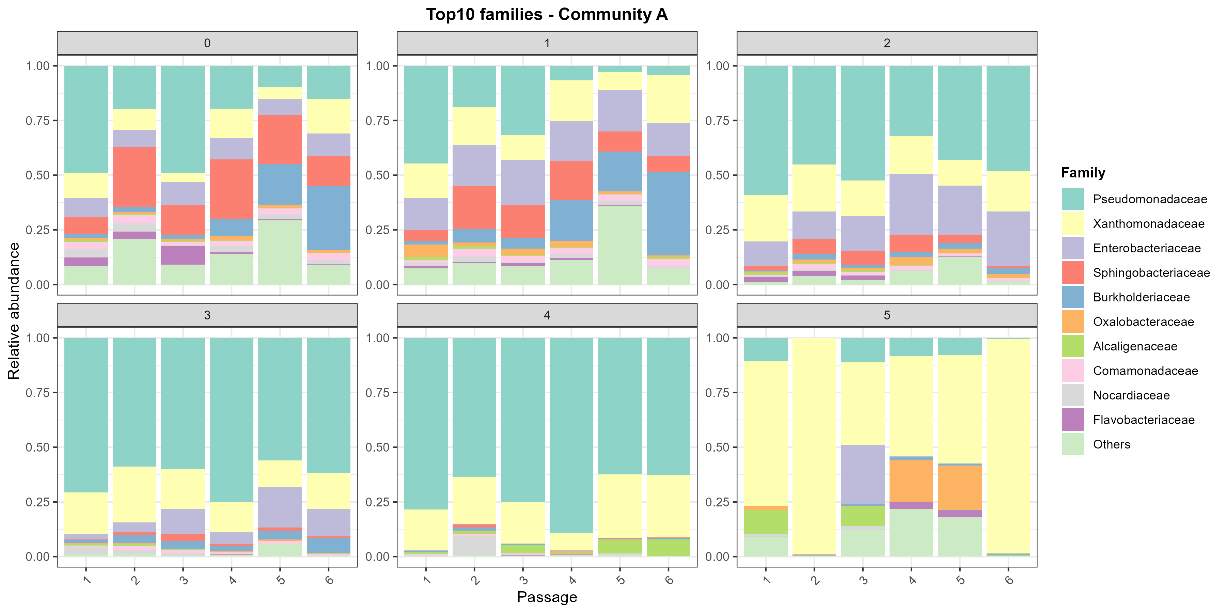

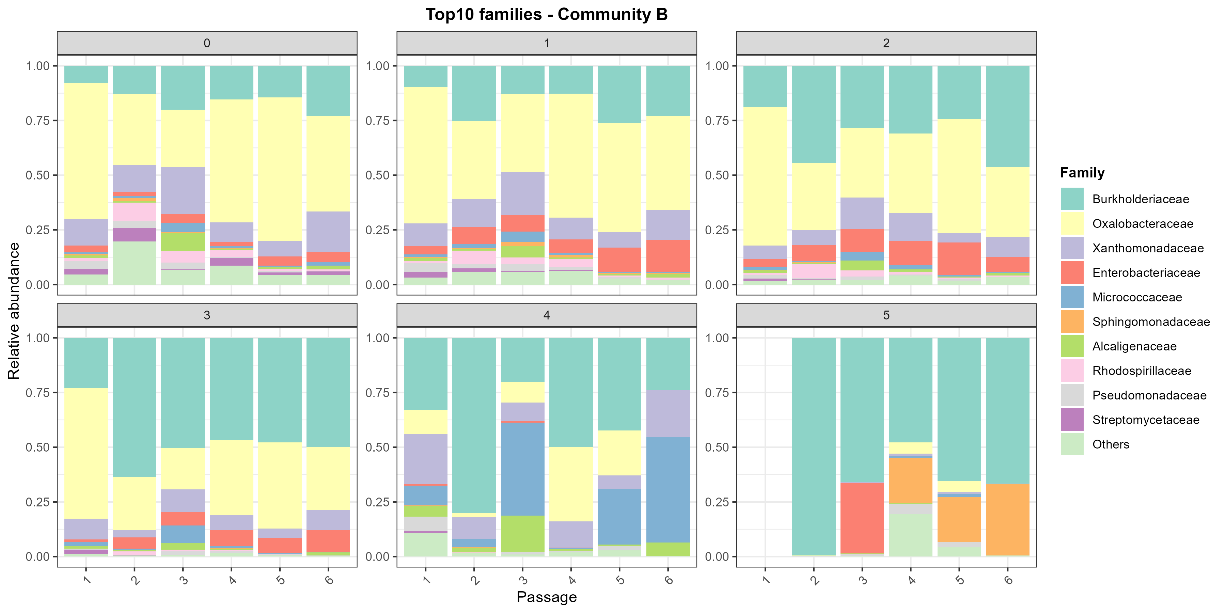

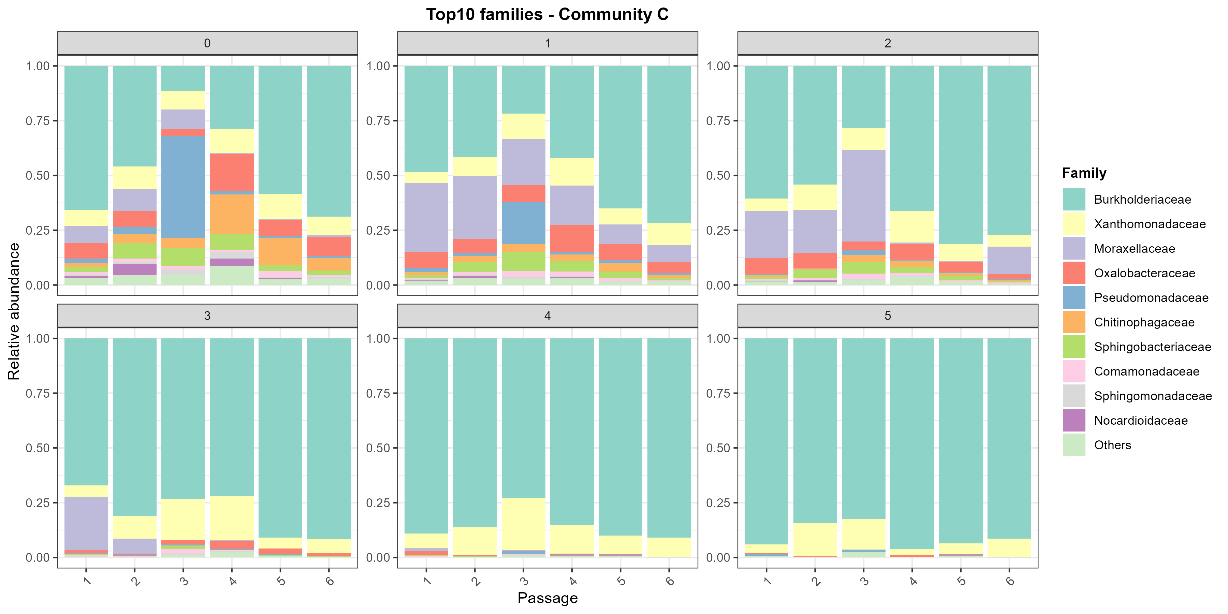

**Figure S1.** Temporal variation in the relative abundance of the ten most abundant bacterial families in tomato plants inoculated with communities A, B, and C. Each panel corresponds to a dilution treatment and shows the distribution of relative abundance values (y-axis) across the different sampling passages (x-axis).

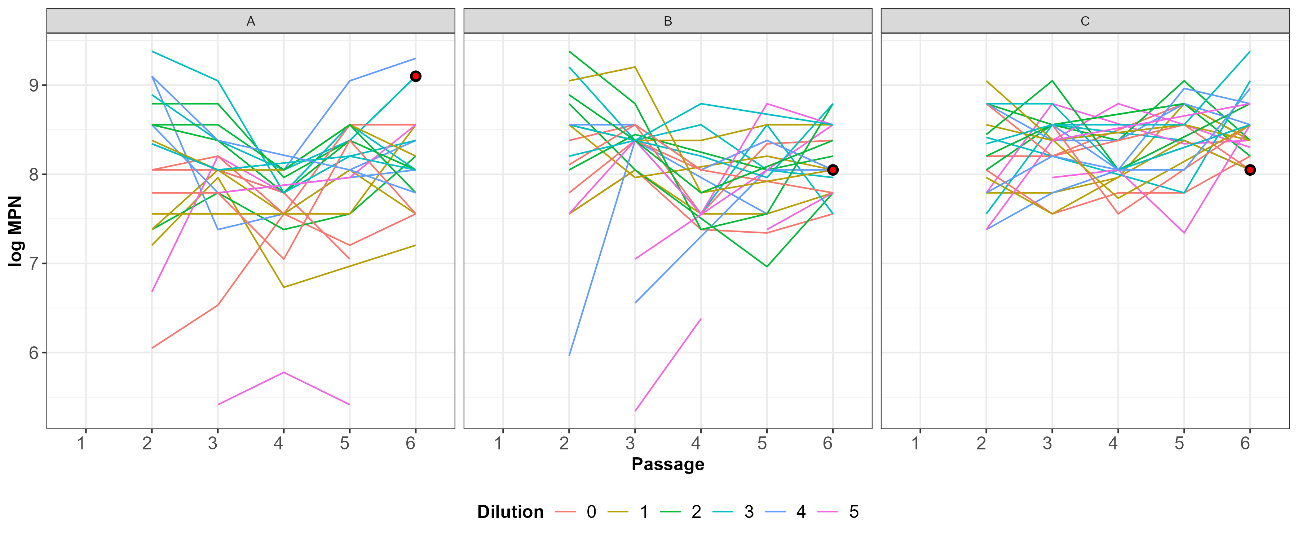

**Figure S2.** Distribution of log-transformed bacterial load as a function of dilution and sampling passage. Each panel corresponds to one inoculated community. Dilution treatments are represented by different colors, while individual experimental trajectories are shown as separate lines. The detection limit for bacterial load was $2.2\times{10}^{5}$ bacteria per plant. The sample selected from each community for subsequent analyses is highlighted with a red circle.

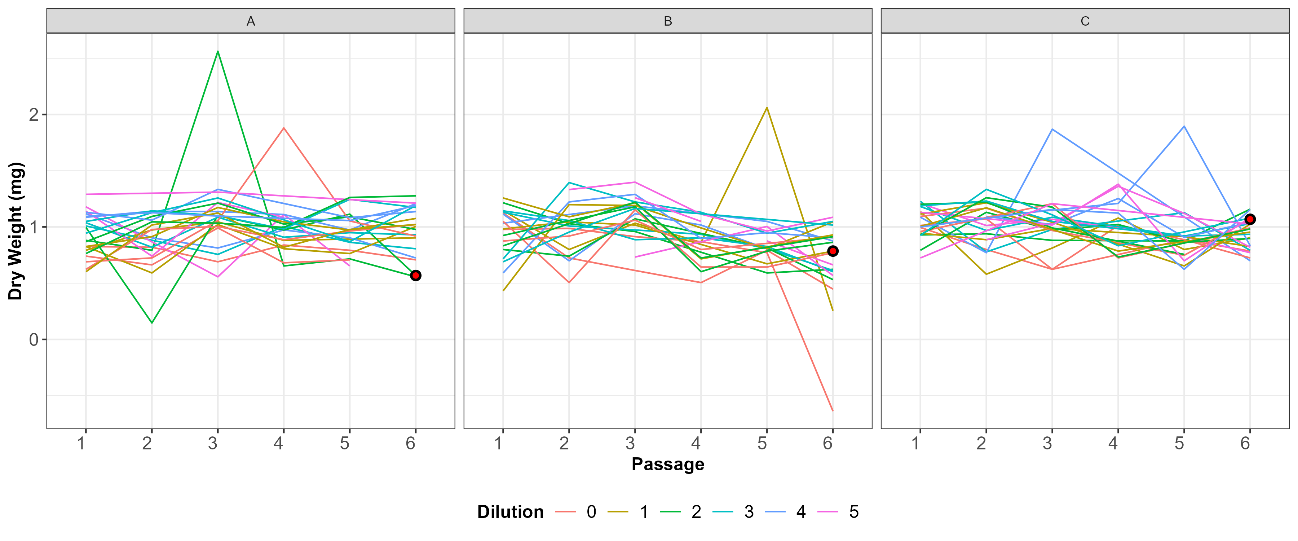

**Figure S3.** Distribution of plant dry weight as a function of dilution and sampling passage. Each panel corresponds to one inoculated community. Dilution treatments are represented by different colors, while individual experimental trajectories are shown as separate lines. The sample selected from each community for subsequent analyses is highlighted with a red circle.
